# Skin resident memory cells generated by the *L. major* centrin gene deleted parasites mediate protective immune response analogous to leishmanization

**DOI:** 10.1101/2021.12.17.472620

**Authors:** Nevien Ismail, Subir Karmakar, Parna Bhattacharya, Telly Sepahpour, Kazuyo Takeda, Shinjiro Hamano, Greg Matlashewski, Abhay R. Satoskar, Sreenivas Gannavaram, Ranadhir Dey, Hira L. Nakhasi

## Abstract

Leishmaniasis is a vector-borne parasitic disease transmitted through the bite of a sand fly with no available vaccine for humans. Recently, we have developed a live attenuated *Leishmania major centrin* gene deleted parasite strain (*LmCen*^*-/-*^) that induced protection against a homologous and heterologous challenges. The protection is mediated by IFNγ secreting CD4^+^ T effector cells and multifunctional T cells, which is analogous to leishmanization. Previously, skin tissue resident memory T cells (TRM cells) were shown to be crucial for host protection in a leishmanization model. In this study, we evaluated generation and function of skin TRM cells following immunization with *LmCen*^*-/-*^ parasites and compared those with leishmanization. In the absence of recoverable *LmCen*^-/-^ parasites, the skin of immunized mice showed functional TRM cells comparable to leishmanized mice. The generation of the skin TRM cells was supported by the induction of cytokines and chemokines essential for their production and survival. Following challenge infection with wild type *L. major*, TRM cells specific to *L. major* were rapidly recruited and proliferated at the site of infection in the immunized mice which was similar to leishmanization. Further, upon challenge, CD4^+^ TRM cells induced higher levels of IFNγ and Granzyme B in the immunized and leishmanized mice than non-immunized mice. Taken together, our studies demonstrate that a genetically modified live attenuated *Leishmania* vaccine generates functional CD4^+^ TRM cells that mediate protection and can be a safer alternative to leishmanization.

## INTRODUCTION

Leishmaniasis is a vector borne neglected tropical disease endemic in tropical and subtropical regions of the world. It is caused by infection with different species of the protozoan parasite *Leishmania* and transmitted by the bites of infected sand fly ^1-4^. Currently there is no approved human vaccine against Leishmaniasis and existing treatment options are sub-optimal because of the development of drug resistance and coinfections with HIV and other endemic diseases ^3,5,6^. Leishmanization, a process by which a small inoculum of *L. major* parasites is injected into the skin to acquire protection against infection was previously used in several countries in the middle east and former Soviet Union ^7^. However, the practice has been discontinued because of safety concerns. Recently, using CRISPR gene editing, we have generated a centrin gene deleted live attenuated *Leishmania major* strain, *LmCen*^*-/-*^ (8). We demonstrated that *LmCen*^*-/-*^ parasites do not cause lesions but have the ability to mount an immunological response that is protective against both cutaneous and visceral leishmaniasis in animal models that mimic human disease ^8,9^. We have shown that both *LmCen*^*-/-*^ immunized and healed mice from primary infection representing leishmanization generated comparable protective immunity against challenge with *LmWT* parasites. The protective immunity was due to multifunctional (IFN-γ^+^ IL-2^+^ TNF-α^+^) CD4^+^T cells as well as IFN-γ secreting CD4^+^ T effector cells (8).

Recent studies have suggested that memory T cells that accumulate in tissues, termed tissue-resident memory T (TRM) cells, play a crucial role in maintaining long-term protective immunity in skin, lungs or any other mucosal organs against viral pathogens and allergens ^10-13^. The CD4^+^ and CD8^+^ TRM cells are identified by the expression of CD69 and CD103 in both mouse and human tissues ^14,15^. In the dermis of mice, the TRM cells are predominately CD69^+^ CD103^-^ while in the epidermis TRM cells are predominantly CD69^+^ CD103^+^. It has been shown that CD69^+^ CD103^+^TRM cells exhibit more effector function compared to CD69^+^ CD103^-14^. The CD4^+^ and CD8^+^ TRM cells can be induced following viral infections at the skin, where they are maintained and capable of rapidly responding to reinfection. TRM cells can also be generated by vaccination, particularly live attenuated viral vaccines appear to be more effective than killed or subunit vaccines for inducing TRM cells ^16-18^.

Previous studies have shown that after the resolution of infection with wild type *L. major* parasites, the skin of healed mice harbors CD4^+^ TRM cells, and the activity of these cells is important for optimal immunity against re-infection with *Leishmania* ^19^. The TRM cells persist in the absence of circulating *Leishmania* specific T cells, rapidly recruit inflammatory monocytes and *Leishmania* specific T effector cells to the site of infection and contribute to protective immunity ^19,20^. In addition, intradermal delivery of a DNA vaccine for *Leishmania* was also found to generate long lasting skin TRM cells that contributed to protective immunity against *L. major* challenge ^21^. TRM cells have been shown to play an important role as the first line of defense in protective immunity in several pathogenic infections. Therefore, TRM cells are an excellent target for vaccine development. In this study we show that intradermal immunization with genetically modified live attenuated *LmCen*^*-/-*^ parasites generate CD4^+^ TRM cells in the skin of C57BL/6 mice. Generation of CD4+ TRM cells in the skin by *LmCen*^*-/-*^ was enabled by the expression of cytokines, chemokine receptors as well as transcription factors as shown in previous studies ^*22*^. The protection induced by *LmCen*^*-/-*^ against infection with virulent *L. major* parasites was due to rapid recruitment, proliferation, induction of Th1 response and cytotoxic response (Granzyme B) by CD4^+^ TRM cells at the site of infection. These observations suggest that intradermal immunization with *LmCen*^*-/-*^ induces a potent protective response comprising CD4^+^ TRM and T effector populations similar to leishmanization and is a safer alternative to leishmanization.

## RESULTS

### Immunization with *LmCen*^-/-^ generates CD4^+^ TRM cells in the skin

Previous studies in leishmanization model have shown that resolution of acute infection with *LmWT* parasites (12-20 weeks post infection), is accompanied by the formation of CD4^+^ TRM cells, that contribute to protective immunity against virulent challenge ^19,20^. To investigate if immunization with *LmCen*^*-/-*^ parasites would similarly lead to the formation of TRM cells, we injected C57BL/6 mice intradermally in the flank skin with 2×10^6^ stationary phase of either *LmCen*^*-/-*^ or *LmWT* parasites and monitored TRM populations in both groups (Fig. 1 a). Parasite load, from the injected flank, and the TRM cell populations in the skin were assessed at 6- and 15-weeks post infection (PI), at the injection site (injected flank) as well as contralateral site (distal flank). We recovered only *LmWT* parasites in the injected flank at 6- but not at 15-weeks PI, suggesting that wild type infection was resolved by 15-weeks PI. On the other hand, *LmCen*^*-/-*^ parasites were not detected neither at 6-nor 15-weeks PI (Fig. 1 b). Next, we evaluated both CD4^+^CD69^+^ as well as CD4^+^CD69^+^CD103^+^ TRM cell populations in the injected and distal flanks at both 6- and 15-weeks post infection/immunization (Fig. 1 c-f). TRM population was identified as CD3^+^CD4^+^CD44^+^CD62L^-^CD69^+^ and CD3^+^CD4^+^CD44^+^CD62L^-^CD69^+^CD103^+^ (Supplementary Fig. 1 A-B). We observed that most of the TRM cells expressing CD69 are also expressing CD103 (Supplementary Fig. 1 a-b). There were very few CD4^+^CD69^+^ as well as CD4^+^CD69^+^CD103^+^ TRM cells in the injected flank of 6 weeks PI in both *LmCen*^*-/-*^ and *LmWT* infected mice (Fig. 1 c-d). However, in the *LmCen*^*-/-*^ immunized mice we observed significantly higher numbers of both CD4^+^CD69^+^ as well as CD4^+^CD69^+^CD103^+^ TRM in the injected flank, at 15-weeks compared to 6-weeks PI or compared to non-immunized mice (Fig. 1 c-d). *LmWT* infected mice (healed mice) also generated significantly higher number of both CD4^+^CD69^+^ as well as CD4^+^CD69^+^CD103^+^ TRM cells at 15-weeks PI compared to non-immunized or 6-weeks PI (Fig. 1 c-d). At the distal flank, there were significantly higher number of both CD4^+^CD69^+^ as well as CD4^+^CD69^+^CD103^+^ (Fig. 1 e-f) TRM cells in the *LmCen*^*-/-*^ immunized animals at 15-weeks of PI compared to 6-weeks PI. On the contrary, distal flank of the *LmWT* group had an increased trend, though not statistically significant, of both CD4^+^CD69^+^ as well as CD4^+^CD69^+^CD103^+^ (Fig. 1 e-f) TRM cells at 15-weeks compared to 6-weeks PI. Interestingly, there was a significant increase of CD4^+^ TRM cells in the distal flank at 15-weeks between *LmCen*^*-/-*^ immunized animals and *LmWT* healed animals (Fig. 1 e-f). We also noted, the frequencies of CD4^+^ TRM cells in the injected and distal flanks are comparable (Fig. 1 c-f), indicating that the TRM cells were present in more or less uniform density throughout the skin of immunized animals. Taken together, these data indicate that immunization with *LmCen*^*-/-*^ parasites generates CD4^+^ TRM cells in the injected as well as distal sites of the skin of mice.

**Fig. 1:**
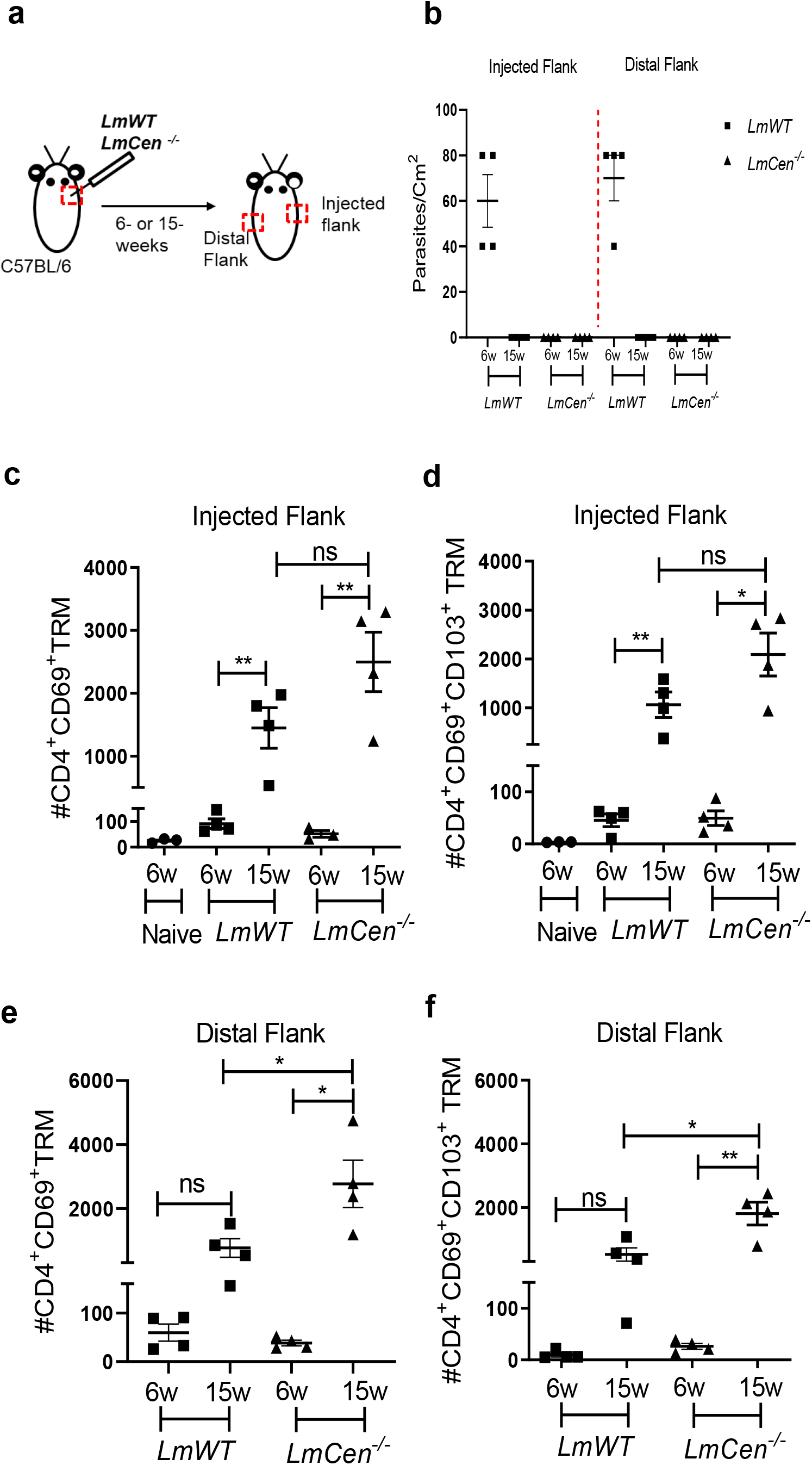
*LmCen*^*-/-*^ immunization generates CD4^+^ TRM cells in the skin. Mice were injected, intradermally, with either *LmCen*^*-/-*^ or *L. major* wild type (*LmWT)* parasites in the right flank and TRM cell population was assessed at 6- and 15-weeks post injections by flowcytometry from both, right (injected flank) and left flank (distal flank) marked by red box. Baseline TRM population was measured in flank skin of non-immunized mice. **a** Schematic plan of injection site and experimental time points. **b** Parasite load per 1cm^2^ of skin, at 6- and 15-weeks post injection, measured by serial dilution. **c-d** TRM cell population in the injected flank **(c)** CD4^+^CD69^+^ TRM cells and **(d)** CD4^+^CD69^+^CD103^+^. **e-f**, TRM cell population in the distal flank **(e)** CD4^+^CD69^+^ TRM cells and **(f)** CD4^+^CD69^+^CD103^+,^ collected from *LmWT* and *LmCen*^*-/-*^ injected mice at 6- and 15-weeks post injection. The Y axis represents number of TRM cells per 10e6 total cells acquired. Results are representative of one of two independent experiments with n=3-4 mice per group. Bars represent the means with SEM in each group. Statistical analysis was performed by unpaired two-tailed t-test (\*\**p*<0.009, \**p*<0.05, ns= not significant).

### Expression profile of cytokines and chemokine receptors supporting TRM cell generation at the injected flank skin

To investigate the immunological milieu that supports TRM cells, we determined the expression of several cytokines and chemokine receptors (AHR, IL15, IL33, CXCR3, CCR8 and TGFB) known to support the formation, survival and homeostasis of TRM cells ^23^. RNA was isolated from the injected flank skin of the *LmCen*^*-/-*^ or *LmWT* injected mice and indicated analytes were assayed by RT-PCR (Fig. 2 a). In the skin of *LmCen*^*-/-*^ immunized mice, the expression of AHR, IL33, CXCR3, CCR8 and TGFB was significantly upregulated at 15-weeks PI compared to 6-weeks PI (Fig. 2 b, d-g). Interestingly, the expression of AHR, IL15, IL33, CXCR3 and CCR8 was significantly higher in the *LmCen*^*-/-*^ immunized group compared to *LmWT* healed mice at 15-weeks PI (Fig. 2 b-f). In the *LmWT* healed group, only AHR, CXCR3 and TGFB were significantly upregulated at 15-weeks compared to 6-weeks PI (Fig. 2 b, e, f). Overall, these results suggest that the cytokine milieu in the skin of *LmCen*^*-/-*^ immunized mice is conducive to support the formation, survival and homeostasis of TRM cells. Further, our results suggest that immunization with *LmCen-/-* parasites results in enhanced cytokine milieu in the skin compared to leishmanization.

**Fig. 2:**
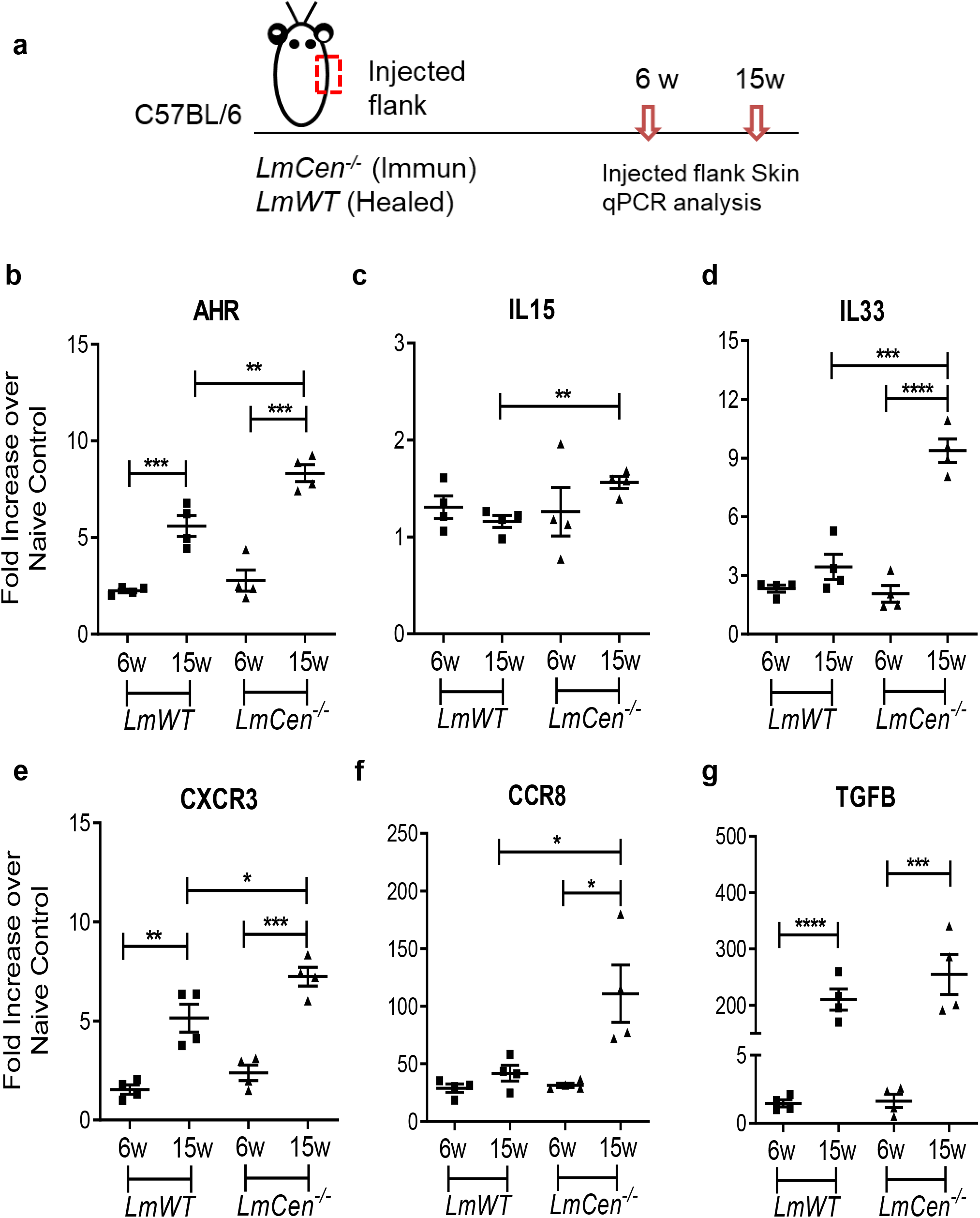
Expression profile of cytokines and chemokine receptors supporting TRM cell generation at the injected flank skin. Mice were injected with *LmCen*^*-/-*^ or *LmWT*, intradermally, in the right flank. The expression profile of indicated genes from the injected flank skin was assessed at 6- and 15-weeks post injection by qPCR. **(a)** Schematic plan of the experimental time points and injection site. **(b-g)** Expression of different transcripts, **(b)** AHR, **(c)** IL15, **(d)** IL33, **(e)** CXCR3, **(f)** CCR8 and **(g)** TGFB at indicated time points. To determine the fold expression of each gene, 2-ΔΔCT method was employed. The data were normalized to GAPDH expression and shown as the fold change relative to age matched naïve mice. Results are representative of one independent experiments, repeated at least 2 times, with total 4 mice per group. Bars represent the means with SEM in each group. Statistical analysis was performed by unpaired two-tailed t test (\**p* < 0.05, \*\**p*<0.009, \*\*\**p*<0.0005).

### *LmWT* challenge leads to rapid accumulation of TRM cells in the skin of immunized or healed mice

To study *Leishmania* specific TRM cell recall response in the skin, non-immunized, *LmCen*^*-/-*^ immunized or healed mice were challenged with *LmWT* parasites. At 48 and 72 hours post-challenge, skin from the challenge site was collected for histology and immune-histochemistry (Fig. 3 a). The challenged flank skin stained with H&E clearly revealed a robust accumulation of cells in *LmCen*^*-/-*^ immunized mice compared to healed or non-immunized mice at 48 hours post challenge (Fig. 3 b). Further, TRM cells in the challenge location were identified by staining for CD3, CD69, and CD103 markers at 48 and 72 hours post challenge (Fig. 3 c). In the immunized mice, challenge infection elicited a rapid increase in the number of CD3^+^CD69^+^CD103^+^ TRM cells as indicated by a strong and focal CD103 staining within 48 hours that persisted till 72 hours (Fig. 3 c). Immunohistochemistry stained section of the challenged site of immunized mice showed more CD69^+^CD103^+^ cells compared to non-immunized or healed mice (Fig. 3 c, inset). In healed mice, the accumulation of TRM cells (CD3^+^CD69^+^CD103^+^) was distinct from non-immunized mice only at 72 hours post-challenge (Fig. 3 c). Non-immunized challenged mice showed little accumulation of TRM cells at the site of challenge which indicates that the rapid accumulation of TRM cells at the site of challenge is a memory specific response to *LmWT* challenge infection.

**Fig. 3:**
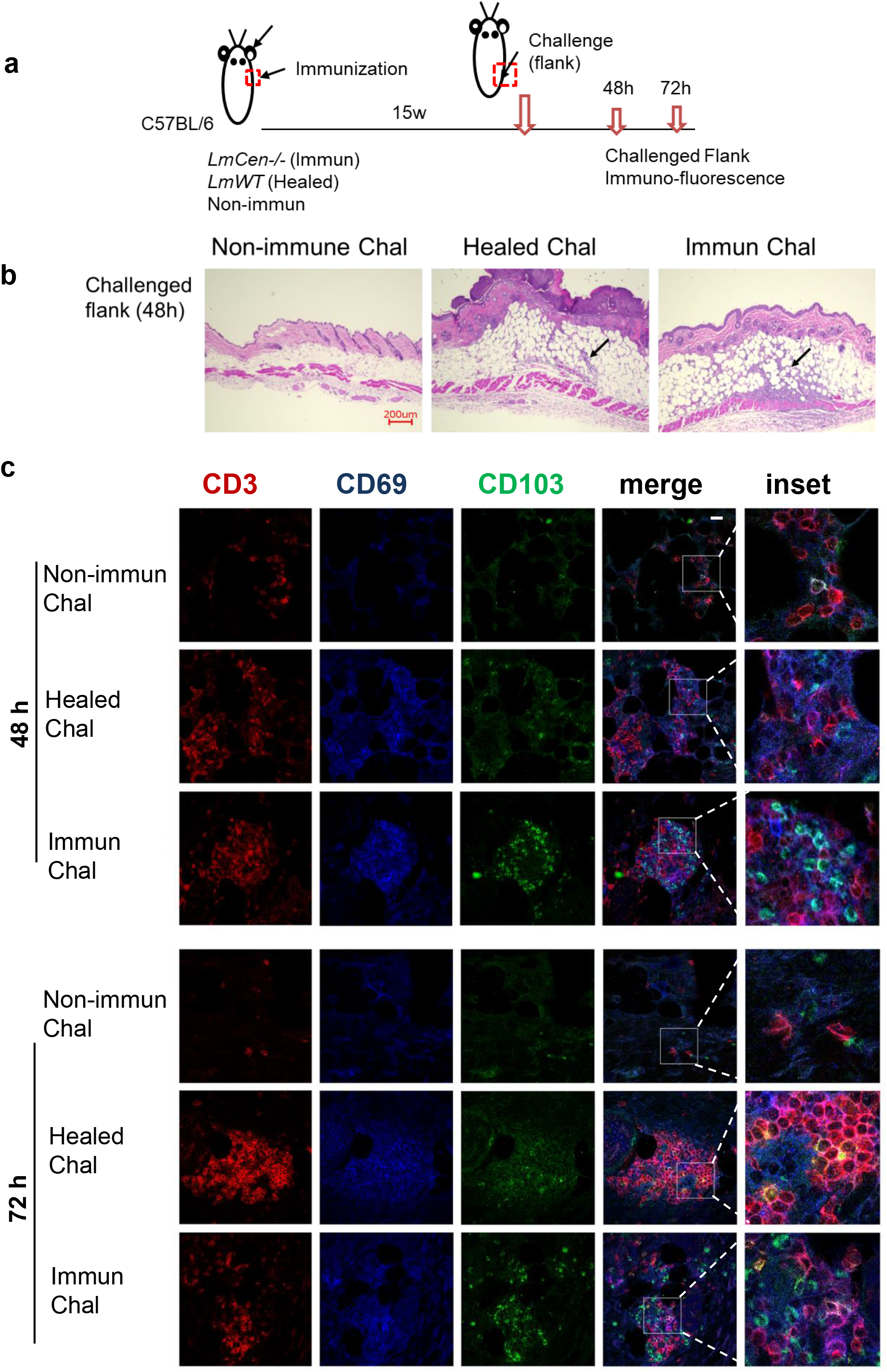
*LmWT* challenge leads to rapid accumulation of TRM cells in the skin of immunized or healed mice. Fifteen weeks Immunized and healed mice as well as non-immunized control mice were challenged in the flank skin with *LmWT* virulent parasite. Skin from the site of challenge was collected at 48h and 72h post challenge. Tissue infiltration and TRM cells were analyzed by immunofluorescence and H&E staining. **(a)** Schematic plan of the experimental time points. **(b)** H&E staining of skin tissue at the site of challenge 48h post challenge. Black arrow indicates cellular infiltration at the challenge site. **(c)** Expression of CD3 (red), CD69 (blue) and CD103 (green) in the flank skin of non-immunized control, healed and immunized mice, at the site of challenge, following 48h and 72h post challenge. Scale bar is 25µm. Results are representative of one independent experiment, repeated 3 times, with 3 mice per group.

### *L. major* specific TRM cells proliferate locally as well as recruit effector T cells after challenge with virulent *LmWT* parasites

It was shown that TRM cells proliferate in situ after an anti-viral recall response ^24^. To assess the proliferative capacity of TRM cells in response to *LmWT* challenge, immunized, healed or non-immunized mice were treated with bromodeoxyuridine (BrdU) for three consecutive days starting on the day of challenge (Fig. 4 a). Seven days post challenge, cells were isolated from the challenged flank skin to detect the proliferating T cells (BrdU^+^ cells) (Supplementary Fig. 2). We observed both immunized and healed mice had significantly higher proportion of BrdU^+^ CD3^+^ T cells compared to the non-immunized challenged group (Fig. 4 b-c). The proportion of BrdU^+^CD3^+^ T cells from immunized-challenged mice was similar to that observed in the healed-challenged group (Fig. 4 b-c). To investigate if the proliferating cells are indeed TRM cells, we measured CD69^+^ as well as skin specific CD69^+^CD103^+^ T cells (Fig. 4 d-f). We found that immunized and healed mice had significantly higher proliferated TRM cells compared to non-immunized challenged mice (Fig. 4 e-f). These results indicate that *LmCen*^*-/-*^ immunization generates skin TRM cells that proliferate in response to challenge with virulent *L. major* parasite. It has been shown in leishmanization model that TRM cells enhance T cell recruitment to the site of challenge ^19^. In our previous study we have shown that, effector T cells may have a role in protection induced by *LmCen*^*-/-*^ immunization ^8^. In the current study, we wanted to investigate if TRM cells generated by *LmCen*^*-/-*^ immunization have any role in the recruitment of *Leishmania* specific T cells from circulation to the site of challenge. Thus, the T cells collected from the spleens of *LmCen*^*-/-*^ immunized mice were stained by CellTrace and injected intravenously (i.v.) into non-immunized, healed or *LmCen*^*-/-*^ immunized mice. The recipient mice were then challenged in the flank skin with *LmWT* parasites (Supplementary Fig. 3 a). To assess T cell recruitment to the site of challenge, we assessed the percentage of CellTrace positive T cells isolated from the challenged flank 48 hours post challenge (Supplementary Fig. 3 b). We observed that upon challenge, the skin of *LmCen*^*-/-*^ immunized mice had significantly higher proportion of CellTrace positive T cells compared to non-immunized challenged mice that lack *L. major* specific skin TRM cells (Supplementary Fig. 3 c). There was no difference in CellTrace positive T cell recruitment between *LmCen*^*-/-*^ immunized and healed mice (Supplementary Fig. 3 c). This suggests that *LmCen*^*-/-*^ specific skin TRM cells enhance the recruitment of T cells from circulation, similar to that observed in leishmanization. Taken together, these results indicate that *LmCen*^*-/-*^ specific skin TRM cells populate the skin at the site of challenge through both *in situ* proliferation and recruitment in addition to effector T cell recruitment.

**Fig. 4:**
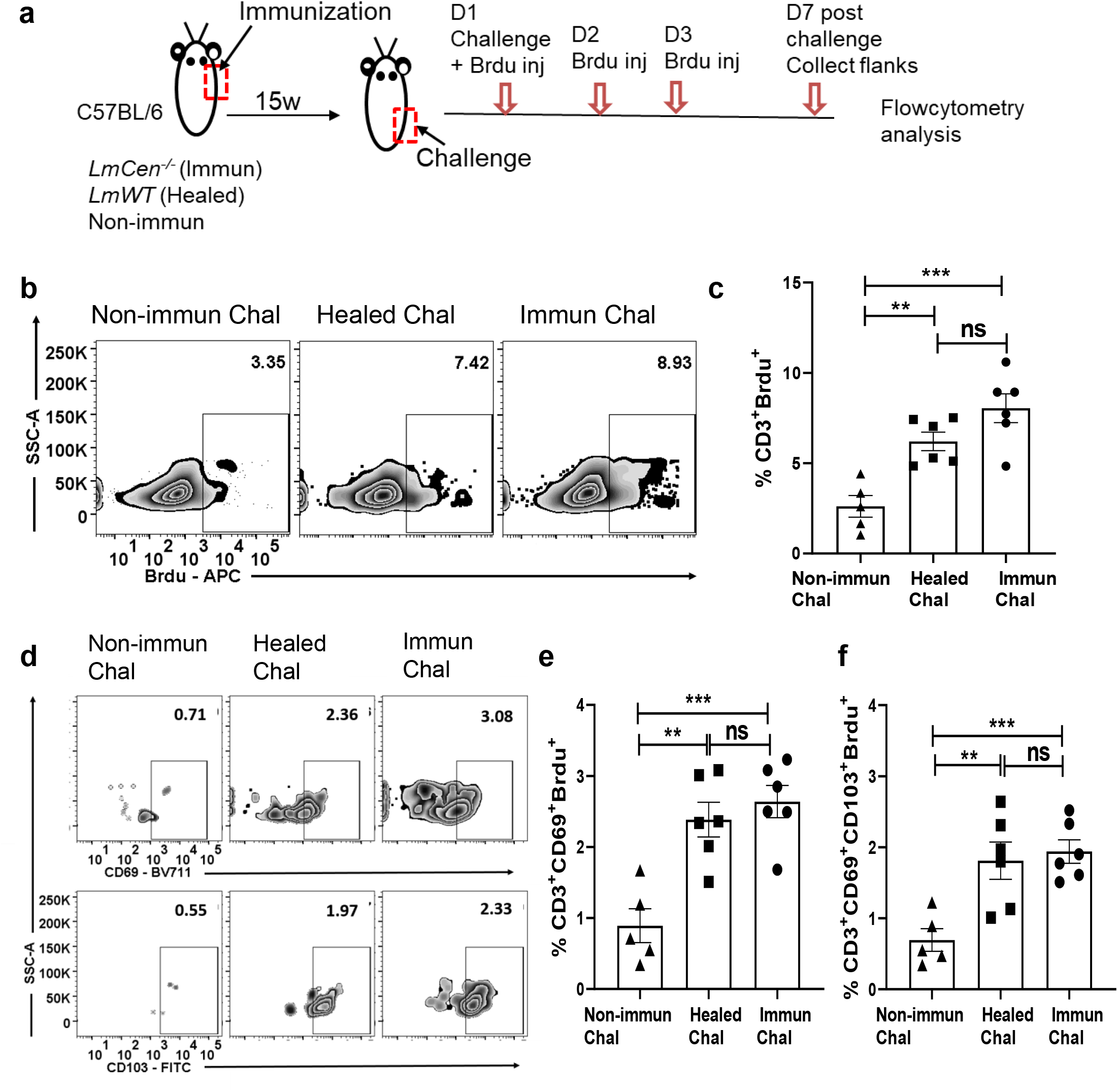
*L. major* specific TRM cells proliferate locally after challenge with virulent *L. major* wild type (*LmWT)* parasites. Fifteen weeks immunized and healed mice were challenged with *LmWT* parasites, in both flanks, and injected with BrdU, as described in the materials and method. Mice were sacrificed and flank skins were collected 7 days post challenge and analyzed for BrdU positive cells **(a)** Schematic plan of the experiment. (**b)** Representative BrdU staining on skin CD3^+^ T cells following 7 days *LmWT* challenge. (**c)** The proportion of skin CD3^+^ T cells with incorporated BrdU in non-immunized (non-Immun Chal), healed (Healed Chal) and immunized (Immun Chal) mice 7 days post *LmWT* challenge. **(d)** Representative BrdU staining on skin CD69^+^ (single positive) and CD69^+^CD103^+^(double positive) TRM cells 7 days post *LmWT* challenge. The numbers represent the percentage of gated population as a ratio of total skin CD3^+^ T cells. **(e)** and **(f)** the proportion of skin CD69^+^ and CD69^+^CD103^+^ TRM cells, respectively, with incorporated BrdU in the Non-immunized, Healed and Immunized mice 7 days post *LmWT* challenge. Y axes in **(b)** and **(d)** represent the percentage of gated population of total skin CD3^+^ T cells. \*\**p*=0.0019, \*\*\**p*=0.0005, ns= not significant. Data shown is combined results from two independent experiments, n=5-6. Results are mean ± SEM, statistical analysis was performed by two tailed unpaired t-test.

### *L. major* specific skin TRM cells produce IFNγ in response to challenge infection

To determine if the TRM cells are capable of secreting IFNγ upon *in situ* reactivation with virulent challenge, without *in vitro* restimulation, we used *IFNγ/Thy1*.*1* knock-in mice, where IFNγ expressing cells could be identified by the surface expression of Thy1.1^23^. Using *IFNγ/Thy1*.*1* knock-in mice increases the sensitivity of detecting IFNγ^+^ cells by flow cytometry. Non-immunized, immunized and healed IFNγ/Thy1.1 knock-in mice were challenged with virulent *LmWT* parasites in the lower flank skin (Fig. 5 a). Five days post challenge, cells were isolated from the challenged site skin and the expression of IFNγ was compared among the groups by flow cytometry (Fig. 5 a-b; Supplementary Fig. 4). We observed that, both CD4^+^CD69^+^ and CD4^+^CD69^+^CD103^+^ TRM cells from immunized mice showed significantly higher expression of IFNγ (represented by Thy1.1 expression) compared to that of non-immunized mice following challenge (Fig 5. c-d). TRM cells from healed mice also had significantly higher expression of IFNγ compared to that of non-immunized mice following challenge (Fig 5. c-d). Importantly, IFNγ expression by TRM cells was comparable between healed and immunized mice (Fig 5. c-d). Taken together these data indicate that *Leishmania* specific skin TRM cells, generated after immunization with *LmCen*^*-/-*^ parasites, can mount a Th1 response upon challenge with virulent parasite.

**Fig. 5:**
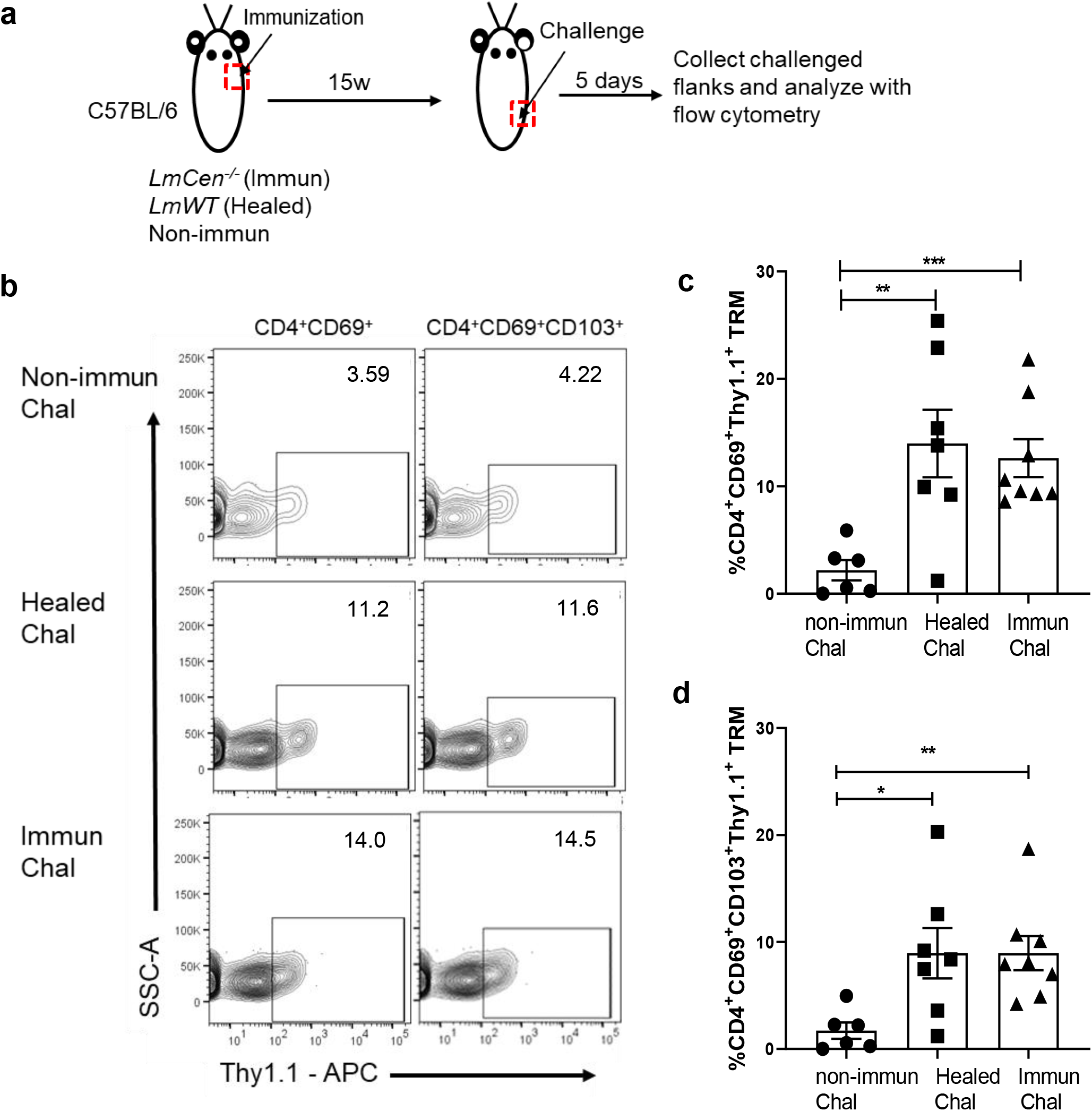
*L. major* specific skin TRM cells produce IFNγ in response to *LmWT* challenge infection. Non-immunized, healed and immunized Ifnγ/Thy1.1 mice were challenged with *LmWT* parasite in the flank skin. T cells from challenged skin were isolated 5-days post challenge and cytokine production was assessed with flow cytometry analysis. **(a)** schematic representation of experiment. **(b)** Graph is representative of Thy1.1 staining on skin CD4^+^ TRM cells (CD3^+^CD4^+^CD44^+^CD62L^-^CD69^+^ and CD3^+^CD4^+^CD44^+^CD62L^-^CD69^+^CD103^+^). **(c, d)** expression of Thy1.1 surface marker, representing IFNγ expression, was measured on **(c)** CD4^+^CD69^+^and **(d)** CD4^+^CD69^+^CD103^+^TRM cells. Y axes represents portion of skin CD4^+^ TRM cells expressing IFNγ as a percentage of parent CD69^+^ TRM population. Data shown is combined results from two independent experiments with n=6-8 mice per group. Bars represent the means with SEM in each group. Statistical analysis was performed by unpaired two-tailed t-test (\*\*\**p*<0.0006 \*\**p*<0.007 \**p*<0.02).

### *Leishmania* specific TRM cells exhibit cytotoxic function by expressing granzyme B

To investigate if the *LmCen-/-* induced TRM cells exhibit cytotoxic function following virulent infection, we challenged non-immunized, immunized and healed mice in the flank skin with *LmWT* parasites (Fig. 6 a). We collected the flank skin 48hours post challenge and stained the tissue with anti-granzyme B antibodies (Fig. 6 a). Immunohistochemistry staining showed that granzyme B expression is higher in the tissue of *LmCen*^*-/-*^ immunized mice compared to healed and non-immunized mice following challenge (Fig. 6 b). The expression of granzyme B in non-immunized mice was minimal (Fig. 6 b). Next, we wanted to investigate if the cells producing granzyme B are indeed TRM cells. Non-immunized, immunized and healed mice were challenged with *LmWT* parasites and granzyme B production by TRM cells was assessed by flowcytometry. We found that significant proportion of CD4^+^CD69^+^ TRM cells, from the skin of *LmCen*^*-/-*^ immunized mice produced granzyme B (∼15%) post challenge, which is significantly higher compared to healed or non-immunized mice (Fig. 6 c). The proportion of granzyme B producing CD4^+^CD69^+^CD103^+^ TRM cells (∼20%) was also significantly higher in the *LmCen*^*-/-*^ immunized mice compared to healed and non-immunized mice following challenge (Fig. 6 d). These results indicate that immunization with *LmCen*^*-/-*^ induced CD4^+^ TRM cells which have a more readily cytotoxic response compared to that of healed mice.

**Fig. 6:**
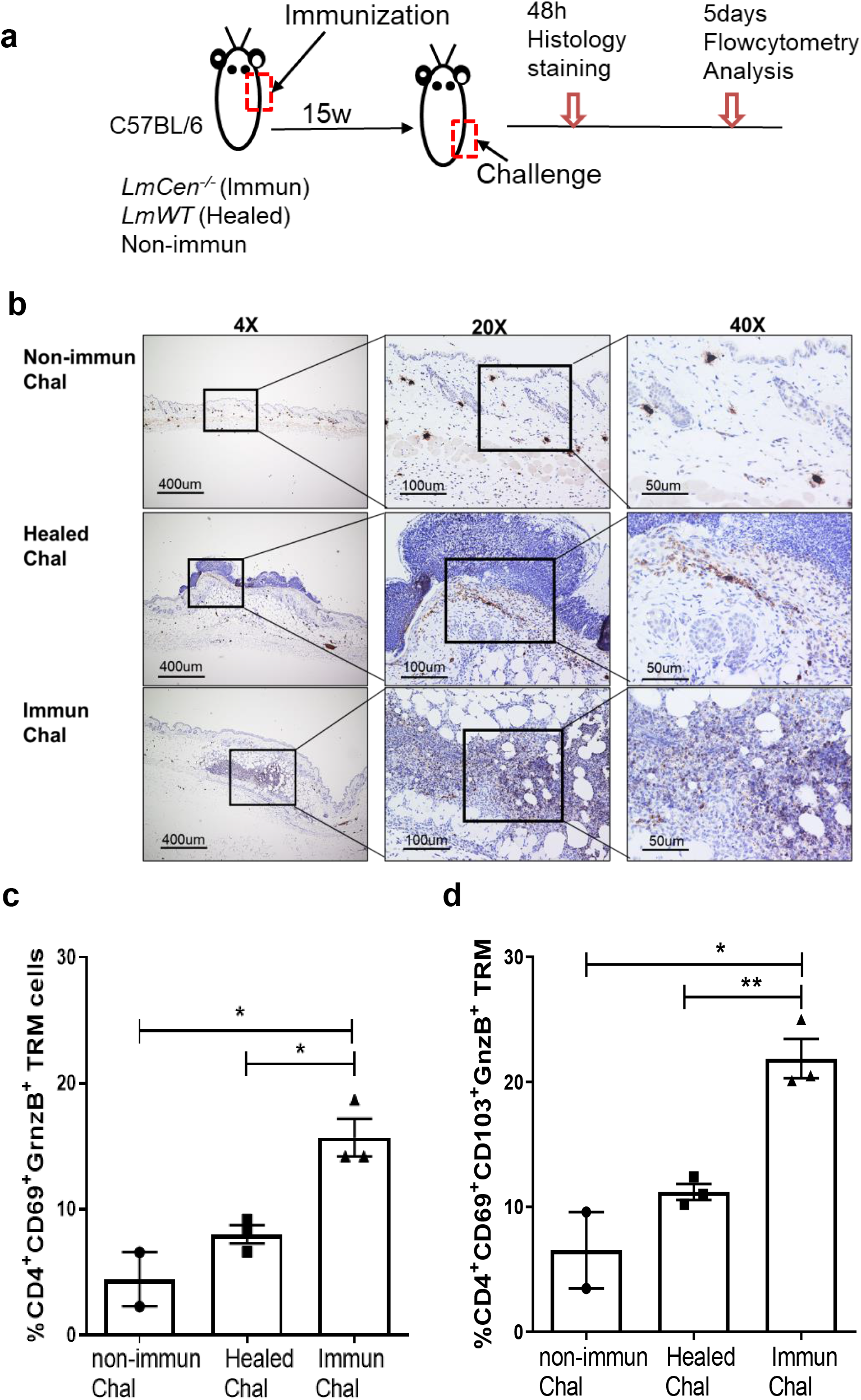
*Leishmania* specific TRM cells produce granzyme B in response to *LmWT* challenge infection. Fifteen weeks Immunized and healed mice; as well as non-immunized control mice were challenged in the flank skin with *LmWT* virulent parasites. **(a)** Schematic diagram of the site of injections. **(b)** Immunohistochemistry of the flank skin, at the challenge site, from 48hrs challenged mice, stained with anti-granzyme B antibodies. **(c-d)** T cells from challenged skin were isolated 5-days post challenge and granzyme B production was assessed with flow cytometry analysis **(c)** portion of CD4 TRM cells, single positive (CD4^+^CD69^+^CD103^+^), expressing granzyme B. **(d)** portion of CD4 TRM cells, double positive (CD4^+^CD69^+^CD103^+^), expressing granzyme B. Results are representative of one of two independent experiments with total 2-4 mice per group. Bars represent the means with SEM in each group. Statistical analysis was performed by unpaired two-tailed t-test (\*\**p*<0.004 \**p*<0.03.).

## DISCUSSION

Skin is the first line of defense against infection with vector-borne pathogens like *Leishmania* parasites. In addition to its physical, chemical and microbiological barriers against pathogens, human skin harbors innate immune cells as well as a combination of resident and recirculating memory T cells with potent effector functions ^14,25^. Upon infection, these cells induce a robust immune response against invading pathogens. In recent years, a T cell lineage termed Tissue Resident Memory (TRM) cells has been identified as the first line of defense against viral infections entering the body at barrier sites, like the skin ^26^. In addition, these cells also play an important role in host immunity against infection with parasites such as *Leishmania* and Malaria ^19,20,27^. Studies in leishmanization models demonstrated that protection upon challenge with *L. major* parasites is mediated by the TRM cells that help recruit heterogenous cell populations including CD4^+^ T effector cells and inflammatory monocytes to the site of infection and mediate parasite control ^19,20^. Prior studies in leishmanization models also showed the critical role of IFN-γ secreting CD4^+^CD44^+^Ly6C^+^ T effector cells in protection against challenge by virtue of their capacity to mount microbicidal activities immediately after challenge ^28,29^. However, since the presence of ‘ready-to-act’ CD4^+^CD44^+^Ly6C^+^ T effector cells requires a persistent infection of *Leishmania* parasites, achieving durable protection through safer vaccination methods makes TRM populations more desirable to target. Such vaccination may be achieved through live attenuated *Leishmania* strains that can persist at low levels in the immunized host yet lack virulence ^8^. Moreover, such vaccines may circumvent the unresolved issues associated with persistent infection of *Leishmania* observed in leishmanization models such as i) indefinite maintenance of T effector populations in presence of concomitant immunity, and ii) potential for T cell exhaustion due to continuous exposure to persisting antigens as discussed previously (28). However, little is known about the role of TRM cells in vaccine immunity in general and in particular in genetically modified live attenuated vaccine candidates including *Leishmania*. In this study, we described live attenuated *LmCen*^*-/-*^ induced skin resident memory T cells and their role in host protection against virulent infection. Recently, we have reported on using CRISPR-Cas9 technology to develop *centrin* gene deleted *Leishmania major* parasites (*LmCen*^*-/-*^). *LmCen*^*-/-*^ parasites showed limited survival in the host and induced protective immunity comparable to leishmanization against both needle and sand fly challenge with a wildtype *L. major* as well as *L. donovani* infection ^8,9^. We also demonstrated that upon challenge with *LmWT* parasites, both *LmCen*^−/−^ immunized and healed mice generated a comparable CD4^+^Ly6C^+^IFN-γ^+^ effector T cells response ^8^, which has been shown to play a role in parasite killing upon re-infection ^30^. In addition to CD4^+^ effector T cells, it was reported that skin resident CD4^+^ TRM cells are generated in response to *Leishmania major* infection after healing and that they play an important role in protection against re-infection along with circulating memory T cells ^19^. In this study, we evaluated the generation and function of skin TRM cells following immunization with *LmCen*^*-/-*^ parasites in mice and compared the response with mice that were healed after a primary *LmWT* infection. TRM cells were identified in our studies by the expression of both CD69 and CD103 as it was previously shown that expression of both the markers is necessary for the optimal formation and survival of TRM cells in the skin ^31^. At 15 weeks post-infection, we could not recover parasites from the injected flank of neither *LmWT* infected mice nor from the *LmCen*^*-/-*^ immunized mice indicating a lack of parasite persistence at this site. The skin of immunized and healed mice sampled at this time point showed a significantly higher population of TRM cells compared to naive control group similar to previous studies ^19,20^. It has been shown that TRM cells are not restricted to the original site of infection, but have the capacity to disseminate throughout the skin ^32,33^. Interestingly, in the *LmCen*^*-/-*^ immunized group, we found that CD4^+^ TRM cells were significantly higher in numbers in the distal sites compared to healed mice suggesting that *LmCen*^-/-^ immunization is efficacious at inducing TRM cell populations.

The role of chemokine receptors in the formation and survival of skin TRM cells has been previously documented ^34,35^. It has been shown that, aryl hydrocarbon receptor (AHR) is required for long term persistence of skin TRM cells ^34,36^. Interestingly *LmCen*^*-/-*^ immunization induced significantly higher expression of AHR compared to healed mice. Similarly, IL15, IL33 and TGFβ are shown to be required for the development and maintenance of TRM cells in the skin following viral (HSV) infection ^31,37,38^. In our study, we observed significantly higher expression of these cytokines in the *LmCen*^*-/-*^ immunized group as well as healed animals compared to non-immunized animals suggesting that immunization with *LmCen*^-/-^ induces an immune milieu that enables the generation of TRM cells, similar to Leishmanization. Future studies will need to address the mechanisms by which these cytokines will help in maintenance of long term TRM cells in the skin of *LmCen*^*-/-*^ immunized mice.

In localized infections, TRM cells are highly protective and modulate host immune response by (i) killing the pathogen-infected cells through direct lysis, (ii) release cytokines that further enhance local recruitment of other innate and adaptive immune cells, and (iii) proliferate *in situ* to maintain a stable population of protective TRM cells ^24,39^. We investigated each of these functions in both immunized and healed mice in response to virulent *LmWT* challenge. Similar to studies in humans, where activated memory CD4^+^T cells, termed CD4^+^ CTL cells, secreted similar amount of granzyme B compared to memory CD8^+^T cells ^40^, CD4^+^ TRM cells from the skin of *LmCen*^*-/-*^ immunized mice produced granzyme B very early on upon challenge. These cells are mainly localized in peripheral tissue like the skin and were found to play an important role in antiviral immunity as studied by others ^40,41^. Parasite specific granzyme B production by human peripheral blood mononuclear cells (PBMC) was found to be a good correlate of protection against *Leishmania*, and could be a biomarker of vaccine induced protection against human leishmaniasis ^2,42^. Our study is the first to show granzyme B production by activated CD4^+^ TRM cells in response to *Leishmania* challenge in mouse model illustrating the direct anti-microbicidal activities of CD4^+^ TRM cells in protection.

We observed pro-inflammatory cytokine, IFNγ, specifically expressed by TRM cells upon challenge with *LmWT* parasites. In addition, as early as 48 hours post challenge, immunized mice recruited an abundant number of immune cells to the site of infection compared to non-immunized challenged mice. These recruited cells included CD69^+^CD103^+^TRM cells that progressively increased in number from 48 to 72 hours post challenge. As shown in previous literature, this increase could be due to proliferation or recruitment from adjacent sites, or from circulation to the site of infection ^24^. Accordingly, we have found that *LmCen*^*-/-*^ specific TRM cells proliferate *in situ* as well as recruit *Leishmania* antigen specific T cells from circulation in response to *LmWT* challenge.

The presence of TRM cells in the lungs has been described as a better surrogate marker, than the memory-specific T cells in the blood, for the protection against viral reinfection in different preclinical studies in mice ^43,44^. Similarly, in non-human primates, the presence of TRM cells against SIV or Ebola virus was essential to control the viral load ^45,46^. As TRM cells are located in the peripheral tissues, early activation and gaining of antimicrobial function is one of the crucial characteristics of these cells. In *Leishmania*, where the rapidity of response is essential for protection against sand fly bite transmission, the presence of skin TRM cells along with effector cells can be the first line of defense keeping the parasites in check until the activation of central memory T cells takes place ^30,47.^ In the current study we observed that within 48 hours post-challenge, CD69^+^CD103^+^ TRM cells accumulated in the challenged site in *LmCen*^*-/-*^ immunized mice, and these cells remained abundant at 72 hours in the infected site. Consistent with the previous studies, the TRM cells secreted IFNγ and granzyme B indicating that their cytotoxic activities maybe important in protection observed in *LmCen*^-/-^ induced immunity.

Overall, our results establish that *LmCen*^*-/-*^ immunization generates CD4^+^ TRM cells in the skin of immunized animals. Upon challenge with wild type *L. major* parasites, these skin TRM cells respond immediately to control the parasites. Moreover, *LmCen*^*-/-*^ immunization induced immunity is comparable with healed mice as measured by effector T cells response ^8^ as well as TRM cells mediated response shown in this study. Since *LmCen*^*-/-*^ parasite are marker free and upon infection do not induce any pathology, it could serve as a safer alternative to leishmanization. Further, since *LmCen*^-/-^ parasites obviate the need for the necessity of persistent infection to maintain protection mediated through CD4^+^ T effector cells and are equally potent in generating CD4^+^ TRM cells compared to leishmanization, they may represent a more practical vaccination strategy with a realistic possibility of gaining approval for clinical use than leishmanization. In conclusion, this preclinical study in animal model further confirms that, *LmCen*^*-/-*^ parasite should be explored as potential *Leishmania* vaccine in future clinical trials.

## MATERIALS & METHODS

### Ethical statement

The animal protocol for this study has been approved by the Institutional Animal Care and Use Committee at the Center for Biologics Evaluation and Research, US FDA (ASP 1995#26). In addition, the animal protocol is in full accordance with “The guide for the care and use of animals as described in the US Public Health Service policy on Humane Care and Use of Laboratory Animals 2015”.

### *Leishmania* strains and culture medium

*L. major* Friedlin (FV9) used in this study were routinely passaged into the footpads of BALB/c mice. Amastigotes isolated from infected lesions were grown in M199 medium and promastigotes were cultured at 27 °C in M199 medium (pH 7.4) supplemented with 10% heat-inactivated fetal bovine serum, 40mM HEPES (pH 7.4), 0.1mM adenine, 5mg L^−1^ hemin, 1mg L^−1^ biotin, 1mg L^−1^ biopterin, 50U ml^−1^ penicillin and 50μg ml^−1^ streptomycin. Cultures were passaged in fresh medium at a 40-fold dilution once a week. The wild type *L. major centrin* gene-deleted *LmCen*^−/−^ (Friedlin strain) promastigotes were cultured as previously described ^8^.

### Mice infection and immunization

Female 6- to 8-wk-old C57BL/6 (Jackson labs) or IFNγ/Thy1.1 knock-in mice ^23^ were immunized or infected, in the upper right flank, with 2 × 10^6^ total stationary phase promastigotes of *LmCen*^*−/−*^ or *LmWT* parasites, by intradermal needle injection, in 10μl PBS. IFNγ/Thy1.1 knock-in mice were provided by C. Weaver (University of Alabama, Birmingham, AL). After 15 weeks of infection/immunization, healed and immunized mice were challenged on the distal flank skin with 2 × 10^6^ total stationary phase *LmWT* promastigotes intradermally by needle inoculation. For measuring the parasite burden from ear tissue, mice were challenged with 2000 wild type *L. major* parasites. Age-matched PBS injected mice were used as a non-immunized control. Parasite burden in the challenged ear was estimated by limiting dilution analysis as previously described ^48^. Briefly, to prepare a single cell suspension from ear tissue, two sheets of ear dermis were separated, deposited in DMEM containing 100U/ml penicillin, 100μg/ml streptomycin, and 0.2mg/ml Liberase CI purified enzyme blend (Roche Diagnostics Corp.), and incubated for 1–2hours at 37°C. Digested tissue was processed in a tissue homogenizer (Medimachine; Becton Dickinson) and filtered through a 70μm cell strainer (Falcon Products). Parasite titrations were performed by serial dilution (1:2 dilutions) of tissue homogenates in 96-well flat-bottom microtiter plates (Corning, Corning, NY) in M199 cell culture media in duplicate and incubated at 26°C without CO_2_ for 7–10 days. The greatest dilution yielding viable parasites was recorded and data are presented the mean parasite dilution ± SEM.

### Flow cytometric analysis

T cells were isolated from the skin using the following protocol: after euthanasia, flanks were shaved and 1cm^2^ of the flank skin was collected, then chopped and incubated in collagenase P (2 mg/mL; Roche Diagnostics) and DNAase in 10% FBS complete RPMI media at 37°C for 120 min. Tissue was then homogenized in MACS C tubes, using gentleMACS Dissociator (Miltenyi Biotec) for 1min. Solution was then strained through 100µm and 70µm nylon strains (Milteny Biotec) to obtain a single-cell suspension. Cells were stained with antibodies, and their expression of phenotypic markers was determined by flowcytometry using BD LSR Fortessa (BD Biosciences) and analyzed with FlowJo (Treestar). We used the following antibodies: anti-mouse CD3 (17A2), CD44 (IM7), Thy-1.1 (HIS51) and Granzyme B (NGZB) from Thermofisher Scientific; anti-mouse CD4 (RM4-5), CD8a (53-6.7), CD62L (MEL-14), CD69 (H1.2F3), CD103 (2E7); anti-BrdU (B24) from BD Biosciences. Live cells were discriminated with a fixable LIVE/DEAD fixable blue dead cell stain (Thermofisher Scientific). Cell number, when indicated, was calculated as an absolute number per 10^6^ total acquired cells.

### Cytokine analysis by RT-PCR

Cytokine expression from mouse tissues (ear and skin) were determined by real-time PCR at the indicated time points. Briefly, total RNA was extracted from flank skin using PureLink RNA Mini kit (Ambion). Aliquots (300ng) of total RNA were reverse transcribed into cDNA by using random hexamers from a high-capacity cDNA reverse transcription kit (Applied Biosystems). TaqMan gene expression Master Mix (Applied Biosystem) was used to determine the cytokine (Table-S1) expression levels in a CFX96 Touch Real-Time System (BioRad, Hercules, CA). The data were analyzed with CFX Manager Software. The expression levels of genes of interest were determined by the 2^-ΔΔCt^ method; samples were normalized to GAPDH expression and determined relative to expression values from naive animals.

### Histological and immunohistochemical staining

Mouse flanks were shaved before tissue harvesting. Flank skin was fixed in fixative solutions (10% buffered formalin phosphate solution). Paraffin-embedded sections were stained with H&E. Immune staining was also done using an anti-GnzB Ab (EPR22645-206) from Abcam. All the histochemical and immunohistochemical staining was done by Histoserv (Gaithersburg, MD). Stained sections were analyzed under a Keyence digital microscope (Keyence Corporation of America). For Immunofluorescence, unfixed mouse skins were embedded in O.C.T. compound embedding medium (Tissue-Tek) and cut 10µm sections for immunohistochemistry. Frozen sections were fixed with cold methanol for 5minutes. Sections were blocked with 10% normal donkey serum and incubated with anti-CD3 (EPR20752), anti-CD69 (H1.2F3) and CD103 (AP-MAB0828) antibodies, from Abcam, overnight at 4°C. After washing with PBS, anti-rat, anti-rabbit, and anti-hamster secondary antibodies conjugated with Alexa Fluor 488, 594 and 647 (Jackson Immuno research) respectively were applied for 1hr at room temperature, washed and followed with Hoechst 33258 nuclear counterstaining, mounted with Fluormont-G. These slides were examined with a Leica SP8 confocal microscope.

### In vivo bromodeoxyuridine (BrdU) treatment

Mice were injected intraperitoneally with 2 mg of BrdU per day, for 3 days, with treatment starting on the day of challenge with *LmWT* parasites. BrdU incorporation was measured with a BrdU flow kit (BD Biosciences), and the proportion of BrdU^+^ cells was measured by flowcytometry and analyzed by flowJo software as indicated in the figure.

### Adoptive Transfer of T cells

For recruitment study, T cells from spleens of 4 weeks *LmCen*^*-/-*^ immunized mice were isolated using mouse Pan T cell isolation kit (Miltenyi Biotec) according to manufacturer protocol. The cells were then stained with Invitrogen CellTrace Far Red Cell Proliferation Kit (Fisher Scientific). The stained cells (30 × 10^6^ /mouse) were transferred intravenously (i.v.) into recipient mice. 24 hours later, recipient mice were then challenged in the flank skin with *L. major WT* parasite (2×10^6^/mouse). Challenged flanks were collected 48 hours post challenge and processed as previously described. Cells were analyzed using flow cytometry.

### Statistical analysis

Statistical analysis of differences between means of groups was determined using a two-tail unpaired t test. All proportional numerical values provided in the text and figure legends were written as the mean ± SEM. All statistical analysis was done in Prism 7.0 (GraphPad). All experiments were performed at least two times, with similar results obtained each time.

## Author contributions

NI, SK, PB, TS and KT conducted experiments, analyzed data; NI, RD and HLN designed the experiments. NI, SH, AS, GM, SG, RD and HLN helped to write the manuscript.

## Data availability statement

The data that support the findings of this study are available from the corresponding author upon reasonable request.

---

**Table S1.**
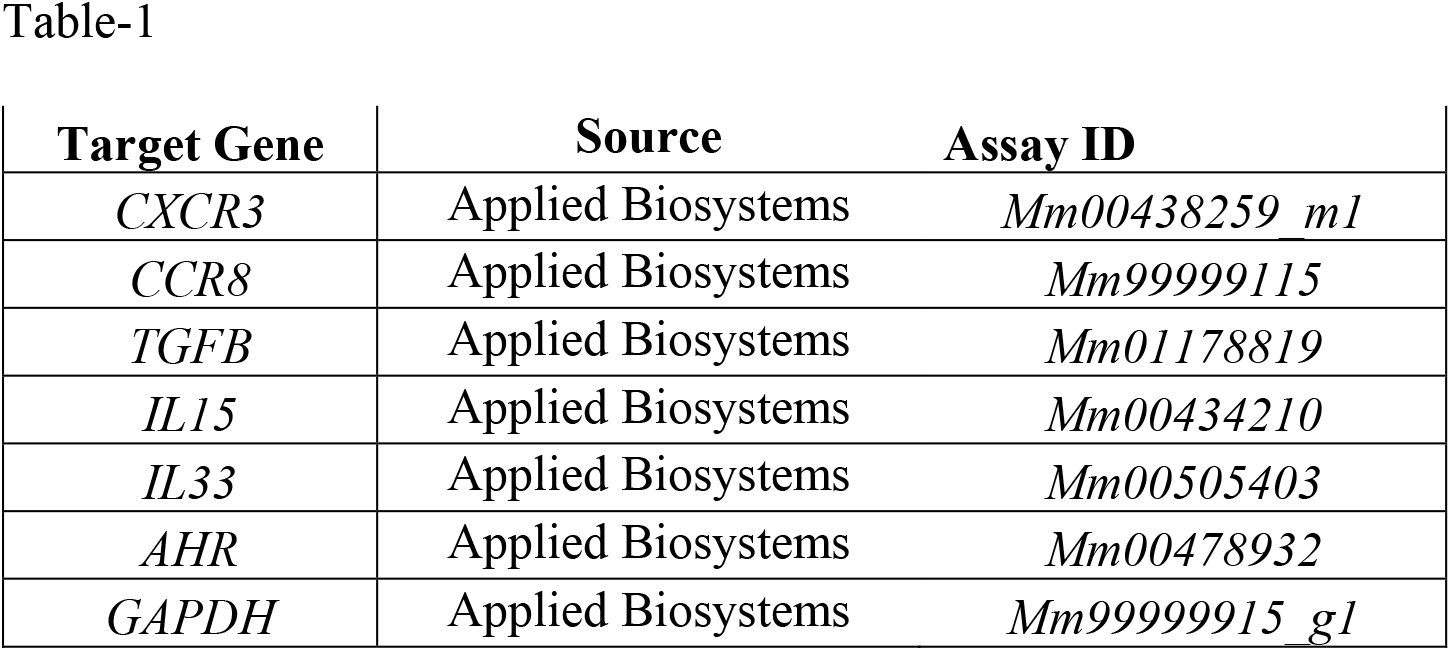
List of genes amplified from RNA extracted from mice skin after injection of *LmCen*^*-/-*^ or *LmWT* parasites Related to main Figure 2.

## Supplemental Figures

**Supplementary Fig 1:**
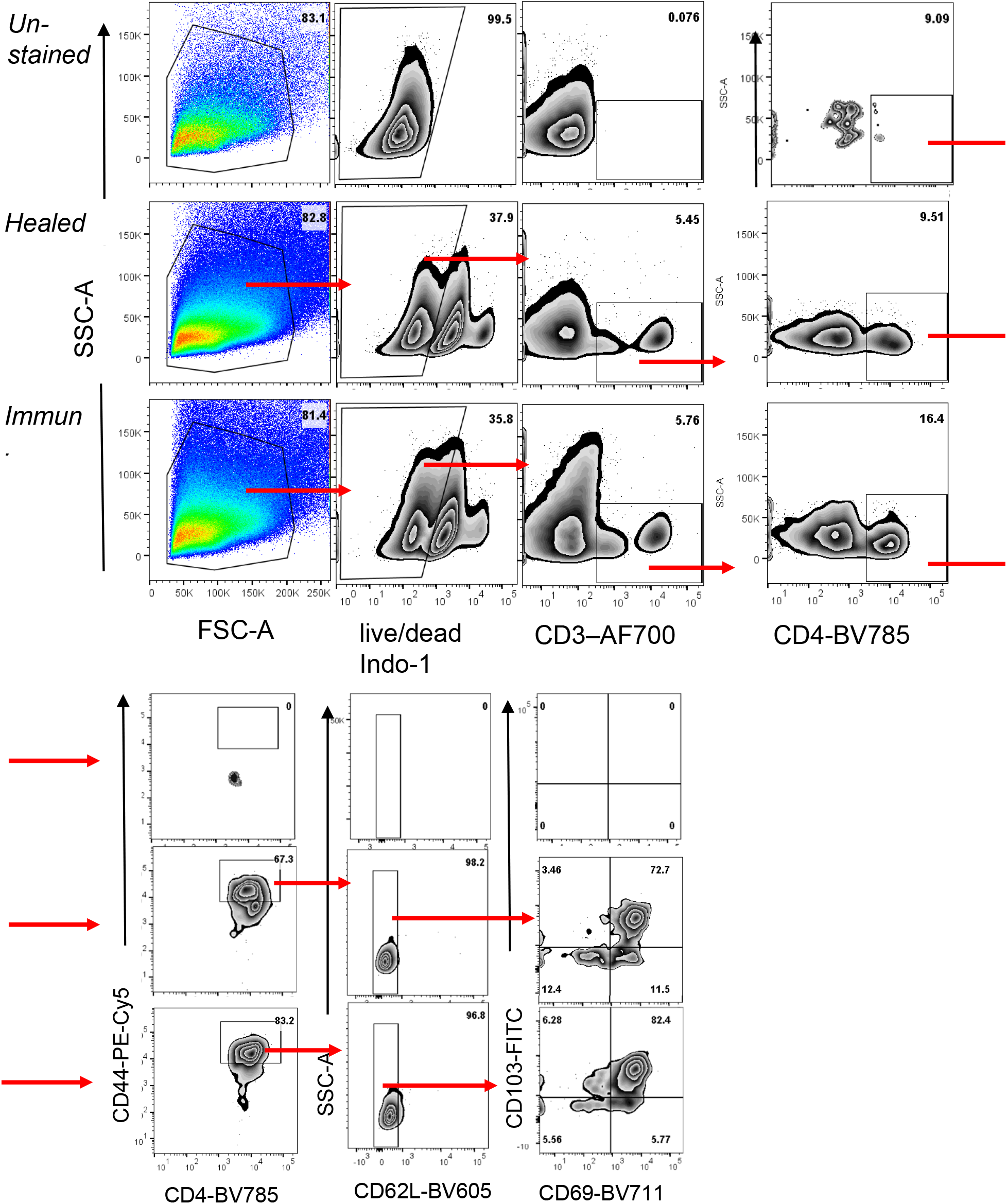
*LmCen*^*-/-*^ immunization generates CD4^+^ TRM cells in the skin. Mice were injected, intradermally, with either *LmCen*^*-/-*^ or *L. major* wild type (*LmWT)*. Baseline TRM population was measured in flank skin of non-immunized mice. **Gating strategy for** **Fig.1 c-f**. Data shown is from 15w post injection time point. Numbers in the quadrants are the proportion of the gated population of parent population

**Supplementary Fig 2:**
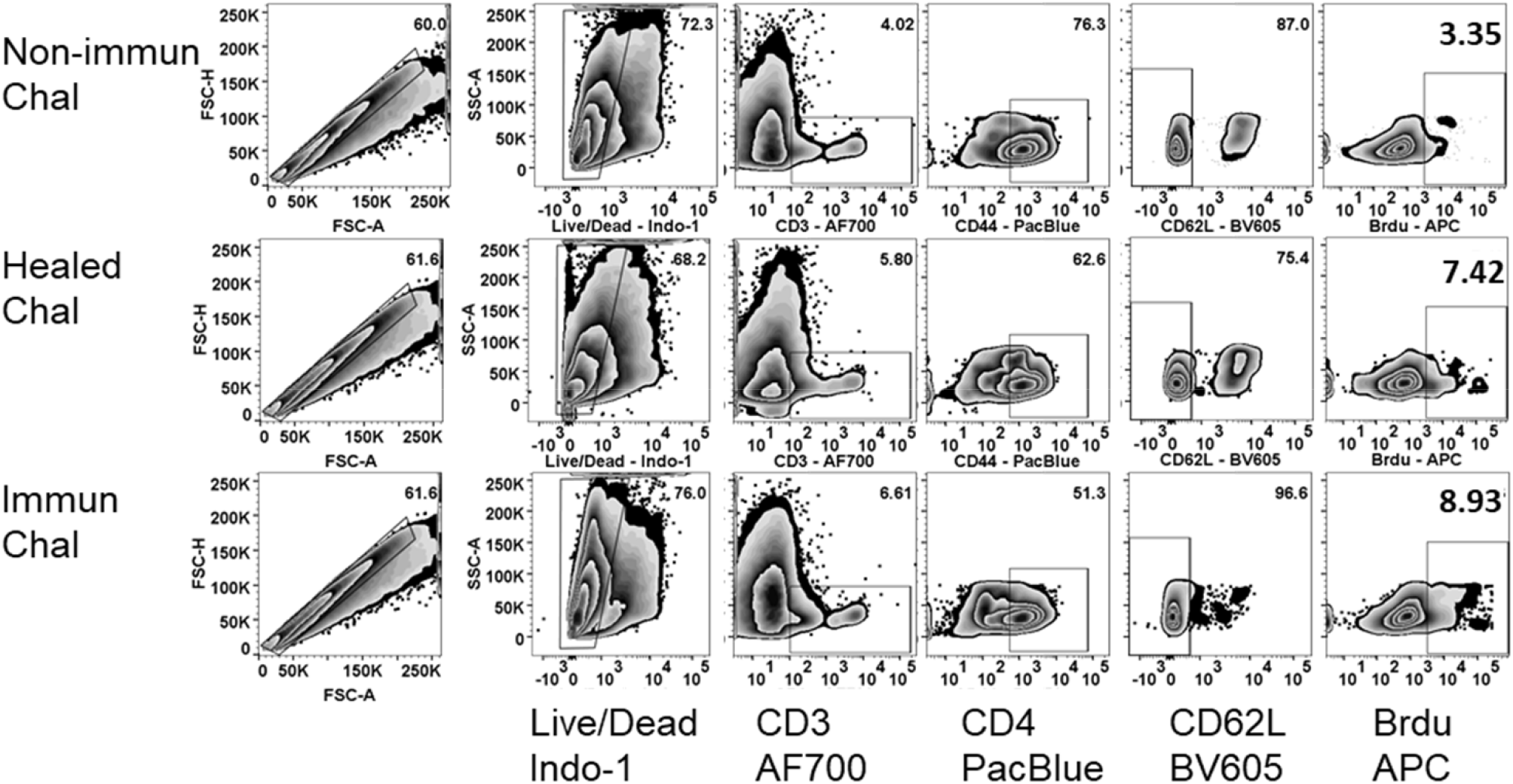
*L. major* specific TRM cells proliferate locally after challenge with virulent *L. major* wild type (*LmWT)* parasites. Fifteen weeks immunized and healed mice were challenged with *LmWT* parasites, in both flanks, and injected with BrdU, as described in the materials and method. Mice were sacrificed and flank skins were collected 7 days post challenge and analyzed for BrdU positive cells. **Gating strategy for** **Fig.4 b**. Numbers in the gate are the percentage of the gated population of the parent population. The zebra graphs are representative of the data combined from 2 independent experiments for 3 groups: non-immunized Challenged (Non-immun Chal) group, n=4; healed challenged (Healed Chal) group, n=6; and Immunized challenged (Immun Chal) group, n=6.

**Supplementary Fig 3:**
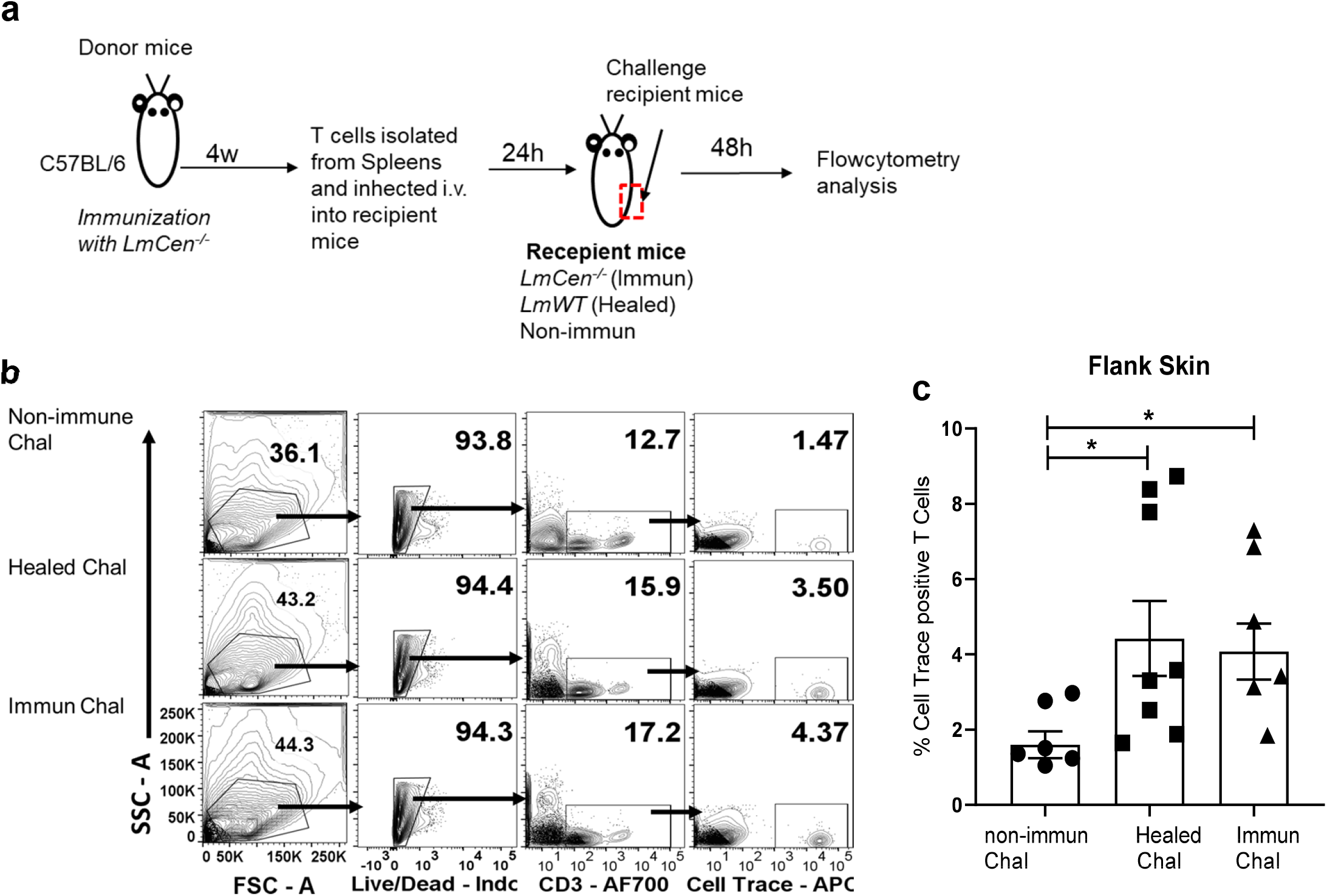
*LmCen*^*-/-*^ specific TRM cells enhance recruitment of circulatory T cell after challenge. T cells from the spleens of *LmCen*^*-/-*^ immunized mice were labeled with Far Red cell trace to track them after transfer. The cells were injected i.v. into naïve, healed or *LmCen*^*-/-*^ immunized mice (15-week post infection). Recipient mice were then challenged with *L. major* WT parasites. 48h post challenge, skin were collected and cell recruitment to the challenge sites were analyzed by flowcytometry analysis for cell trace positive cells. (a) Experimental diagram. (b) Gating strategy representative flow plots of Far Red^+^ CD3^+^ T cells from the flank skin challenged mice. Numbers in the quadrant represents the percentage of the gated population of parent population (c) Frequency of transferred *LmCen*^*-/-*^ immune CD3^+^ T cells in the flank skin of non-immune challenged, healed challenged or immune challenged mice 48h post challenge. Y axes represents the percentage of gated population of total skin CD3^+^ T cells. \**p*<0.03. Data shown is combined results from two independent experiments, n=6-8. Results are mean ± SEM, statistical analysis was performed by tow tailed unpaired t-test

**Supplementary Fig 4:**
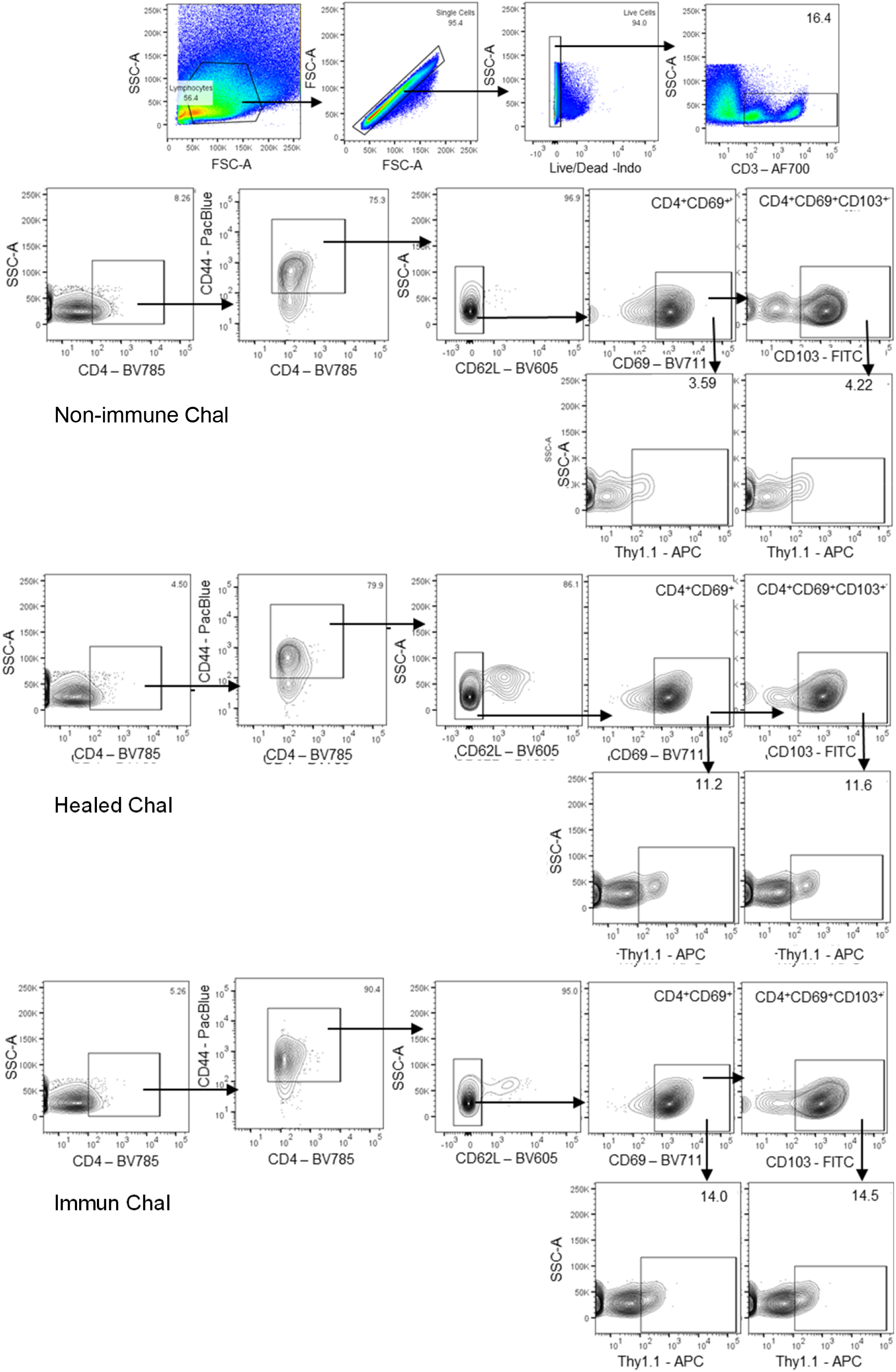
*L. major* specific skin TRM cells produce IFNγ in response to *LmWT* challenge infection. Non-immunized, healed and immunized Ifnγ/Thy1.1 mice were challenged with *LmWT* parasite in the flank skin. T cells from challenged skin were isolated 5-days post challenge and cytokine production was assessed with flow cytometry analysis. Gating strategy shown is representative of samples acquired. Numbers in the quadrants are the proportion of the gated population of parent population

